# Validated method for automated glioma diagnosis from GFAP immunohistological images: a complete pipeline

**DOI:** 10.1101/2022.01.09.474689

**Authors:** A. Campo, F. Fernández-Flores, M. Pumarola

## Abstract

**Background and objective:** Glial fibrillar acid protein is a common marker for brain tumor because of its particular rearrangement during tumor development. It is commonly used in manually histological glioma detection and grading. An automatic pipeline for tumor diagnosis based on GFAP is proposed in the present manuscript for detecting and grading canine brain glioma in stages III and IV.

**Methods:** The study was performed on canine brain tumor stages III and IV as well as healthy tissue immunohistochemically stained for gliofibrillar astroglial protein. Four stereological indexes were developed using the area of the image as reference unit: density of glioma protein, density of neuropil, density of astrocytes and the glioma nuclei number density. Images of the slides were subset for image analysis (n=1415) and indexed. The stereological indexes of each subset constituted an array of data describing the tumor phase of the subset. A 5% of these arrays were used as training set for decision tree classification with PCA. The other arrays were further classified in a supervised approach. ANOVA and PCA analysis were applied to the indexes.

**Results:** The final pipeline is able to detect brain tumor and to grade it automatically. Added to it, the role the neuropil during tumor development has been quantified for the first time. While astroglial cells tend to disappear, glioma cells invade all the tumor area almost to a saturation in stage III before reducing the density in stage IV. The density of the neuropil is reduced during the tumour growth.

**Conclusions:** The method validated ere allows the automated diagnosis and grading of glioma in dogs. This method opens the research of the role of the neuropil in tumor development.

**Graphical abstract:** 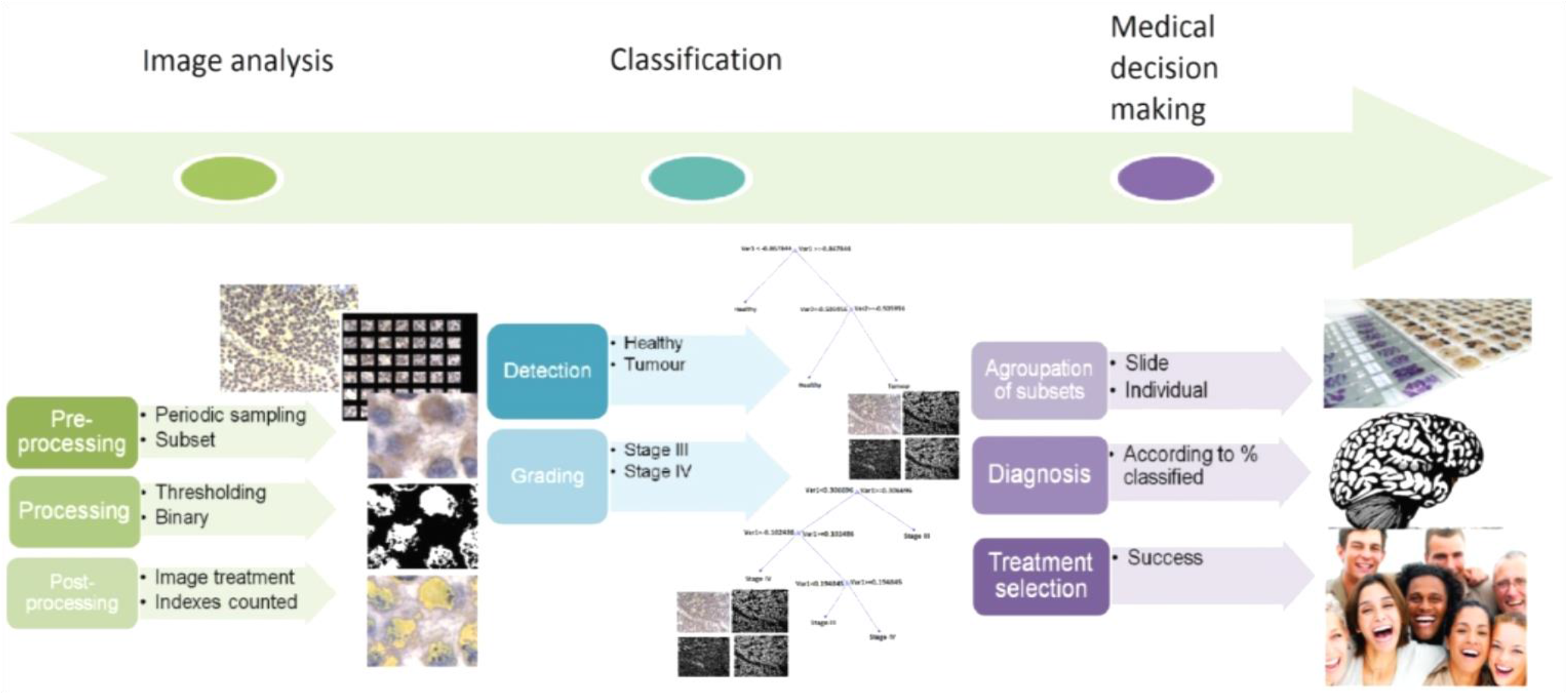

## 1 Introduction

Objective assistance in critical decisions can increase the efficiency of the diagnosis and prognosis of many sickness, including cancer [1–7]. Many tumor classification procedures use machine learning tasks for cancer diagnosis are based on fMRI and DNA microarrays [8–14]. From those, only few machine-learning methods have been applied to histological image analyses. Previous studies from histological analysis included colon cancer [7], breast cancer [15] and prostate cancer [16,17] for quantification of key structures such as cell number. Image analysis packages [17,18] automate this quantification recognising the morphology and color of these features and distinguishing the regions of interest from the background or from other features with the same color. Image segmentation algorithms such as thresholding, edge detection and region growing [3], are able to distinguish tumor features from classically stained histological sections and to classify them depending on the tumor malignancy grade [2–4,6,19–21].

However, the quantification from two dimensional slice of tissue produces indexes that do not take into account the volumetric size, shape, or orientation of particles, and therefore these indexes are biased [22]. In order to proceed with an unbiased estimation of the features of interest, stereological algorithms correct the error thus providing with quantification indexes related to the volumetric distribution of the indicators. These algorithms allow the three-dimensional interpretation of two-dimensional cross sections of materials or tissues [23]. Added to this, the quantification by stereological unbiased indexes can increase the accuracy of the diagnosis and with it the possibilities of success in treatment selection [24–30].

In the present article we propose a diagnosis of glioma, brain tumor, based on segmentation of the extracellular matrix (ECM) which in turn is composed by glial fibrillar acid protein (GFAP). The high mitotic activity of the astrocytic neoplastic cells in the brain induce an asymmetric distribution of the GFAP filaments in the cytoplasm of those cells [31]. Immunohistochemical (IHC) techniques allow the specific staining of cancer markers to ease the slide evaluation process [32–34] and it has been developed for GFAP [35]. The features of interest in our analysis are the astrocyte GFAP (AG), neuropil GFAP (NG), glioma GFAP (GP) and glioma nuclei (GN).

After the segmentation we proceed with a stereology-based indexation using uniform periodic sampling and classification by decision tree learning approach. Following the World Health Organization (WHO) tumor grading procedures, diagnosis by classification is divided in two basic steps: tumor detection and tumor grading. We create a training set of images and we evaluate other slides after the classification by PCA.

In our approach, canine glioma samples were used as model for human glioma because of the similar incidences to that of humans [36] and common neuroimaging, clinical course and immunomodulation features [37].

## 2 Material and methods

### 2.1 Cases

Nine spontaneous canine gliomas were selected from the database of the Veterinary Neuropathology group of the *Universitat Autònoma de Barcelona*, including seven Anaplastic Oligodendroglioma (WHO grade III) and two Glioblastomas (WHO grade IV). The presumptive diagnosis of glioma was made by a Board-certified neurologist from the neurology service of the Veterinary Teaching Hospital of the *Fundació Hopital Clínic Veterinari* of the same university. This diagnosis is based on clinical criteria, magnetic resonance imaging features, and cerebrospinal fluid analysis. The diagnosis was confirmed after necropsy and by histopathological analysis. The owners provided written consent for all the analysis and the publication of the results.

### 2.2 Histological evaluation

Representative tissue samples were collected from all the organs including the brain, fixed in 10% formalin, dehydrated in ethanol and embedded in paraffin. Sections of 5μm were obtained from paraffin-embedded tissues, and stained with haematoxylin and eosin (HE) for microscopic evaluation and diagnosis. The morphological diagnosis and grading of canine gliomas was performed by the pathologists based on the specific histopathological features described in human gliomas [38].

### 2.3 Immunohistochemistry

The IHC glial cell marker used was a rabbit anti-GFAP protein polyclonal antibody (Dako, ZO334). For the antigen retrieval, sections were heated during 20 min in a water bath with 10 mM citrate buffer at pH 6.0. Then, samples were cooled for 30 min at room temperature and rinsed in phosphate-buffer saline (PBS). Previously, sections were treated 35 min with 3% peroxidase to block endogenous peroxidase activity. Non-specific binding was blocked by normal goat serum 30% diluted with PBS during 1h at room temperature. Samples were incubated overnight at 4 ºC. Sections were rinsed with PBS and incubated for 40 min with a labelled polymer according to the manufacturer’s instructions. After, an anti-rabbit EnVision™ kit (catalogue number K011, Dako) was used to label the primary antibodies. Staining was completed by a 10 min incubation with 3,3′-diaminobenzidine (DAB) and counterstaining in haematoxylin for 3 s. The positive control tissue was normal brain including cortical grey matter and subcortical white matter. In all IHC procedures, negative controls were used omitting the primary antibody and no immuno-reaction was observed.

### 2.4 Image intake

Images were taken with a Leica DM6000 camera attached to a Leica DM4 microscope. The output image resulted in a complete section of tissue and it is referred here as original image. Four to five images per individual were taken from different areas of the tumor, by which at least 95% of the image was covered.

### 2.5 Analysis pipeline

For analysing the images, a software was developed using MATLAB 2016a following the standard pipeline: pre-processing, processing and post-processing (Table 1).

**Table 1:**
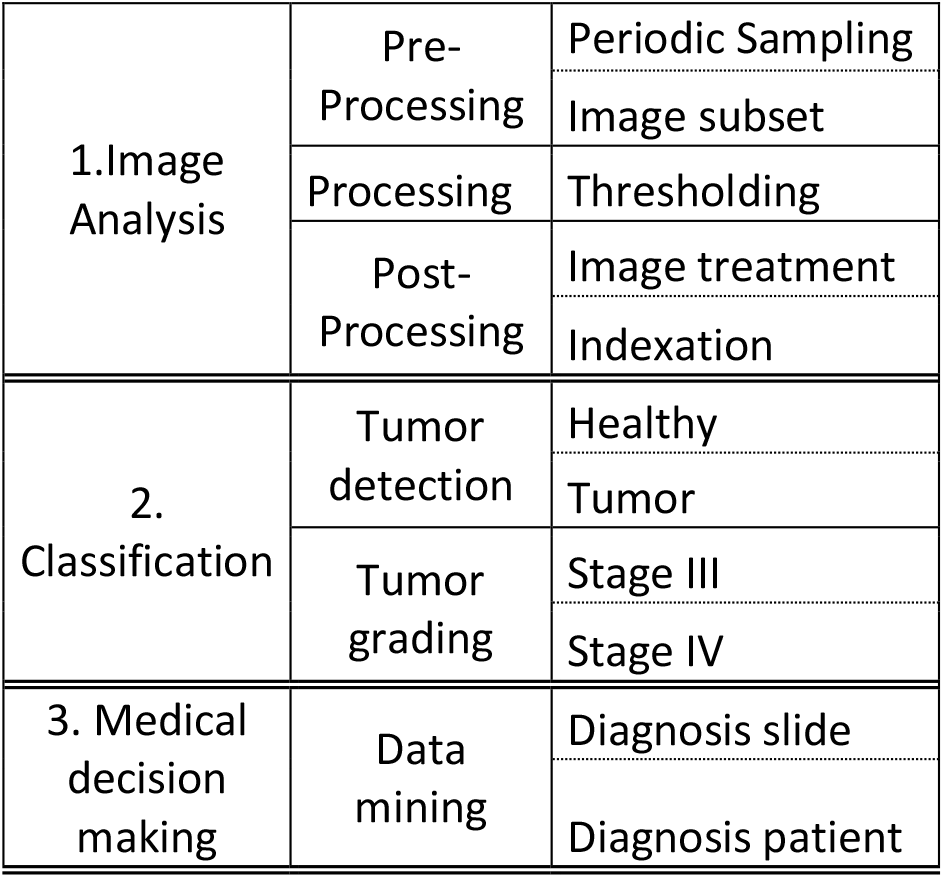
Complete working pipeline: Image analysis, tumor classification and medical decision-making. Image analysis is divided in pre-processing, processing and post-processing, while classification is performed first for detection and later for grading. The results of the classification are further grouped by image and by individual. The final diagnosis for supporting the medical decision-making is established with the previous classification per individual.

#### 2.5.1 Pre-processing: image sampling

Three methods of image sampling were compared by studying the reduction of the standard error [30]. These methods are complete, random and periodic sampling. In the complete method, the original image was analysed and here we will refer it as complete sampling method or CS (Figure 1A). The same analysis was repeated sub-setting the complete image. The image is sub-sampled in smaller rectangles or subsets that allow an array of sub-images per individual. The informative indexes were analysed within those images to perform the statistical analysis with higher accuracy and reduction of the standard error. Image sampling produced subsets of 250×200 pixels. The size of the subsets was chosen to include representative cell features of interest with grading purposes. Each subset has a start point or origin (x,y) placed in the top left corner. Those starting points were declared following two different methods: random and periodic.

**Figure 1:**
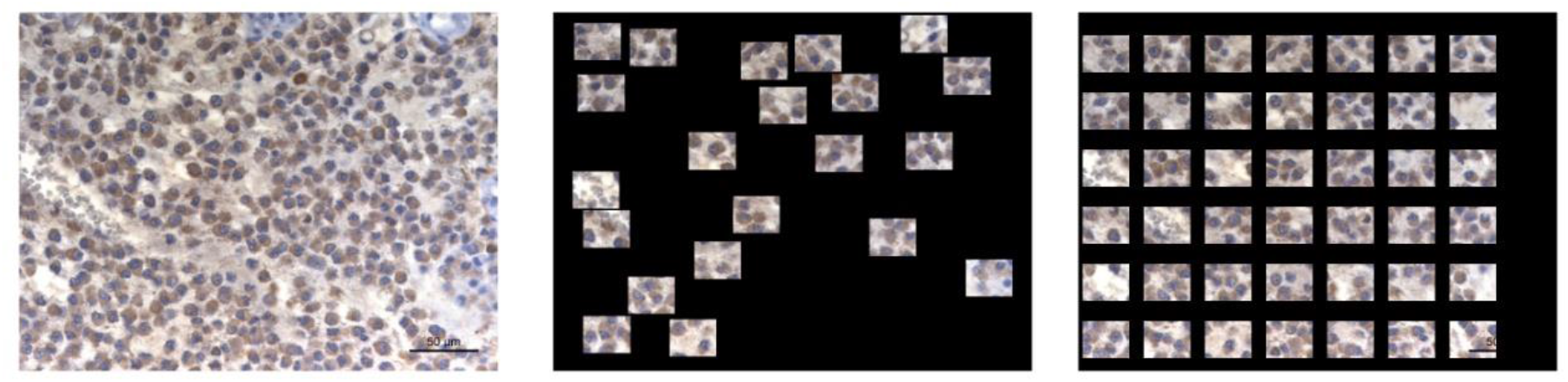
Image analysis method A: Complete analysis of the histological section. B: Random sampling of the histological image. C:Uniform Periodic Sampling of the histological image. The analysis took place on the image areas, while the black areas where not taken into account.

The first method consisted in selecting random starting points in the full image and it is called here random sampling method or RS (Figure 1B). There were starting points overlapping already taken areas and were avoided by the program.

The last method for analysing the image is the periodic sampling. In this method there is a unique starting point as origin selected from the first 250×200 pixels in the top left side of the image. Two periods, one for each axis, x and y, are established from this point thus creating an imaginary grid with squares never smaller than 250×200 pixels. The crossing points of this grid will become the origin of the subsets (Figure 1C). The subsets were selected in a periodic manner and no overlapping occurred. This method is therefore named periodic sampling or PS and achieves the instruction in Weibel’s protocol for analysis of histological images [30].

In order to obtain only subsets fitting in the original image in the periodic sampling, their number (*n*) was limited using the following relation:

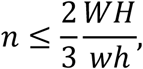

where *H* is the height of the original image, *W* the width of the original image, *h* the height of the subset (200 pixels) and *w* the width of the subset (250 pixels). Only the equality was considered.

The period *p*_*x*_ of horizontal sampling and *p*_*y*_ of vertical sampling were taken as

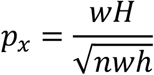

and

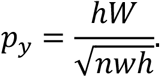

The origin *(x*_*0*_,*y*_*0*_) for sampling is defined by

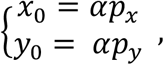

where α is a random number taking values between 0 and 1.

More than 40 subsets per image were obtained with both systems, random (total subsets = 658) and periodic (total subsets = 1415). The features quantified from CS analysis of the image have provided four values per individual. In contrast, the features indexed for each of the subsets provided more than 40 values per individual. Therefore, each ID could have average and standard deviation for each index. This standard deviation will be taken into account for a multilevel statistical analysis.

#### 2.5.2 Processing: Segmentation

Our team has segmented 4 basic structures for glioma grading: astrocyte GFAP (AG), neuropil GFAP (NG), glioma GFAP (GP) and glioma nuclei (GN).

The protein synthesized by the glial cells (GFAP) was immuno-stained with the previously described antibody. Three different stain intensities were detected according to the proteomic structure of GFAP: synthesized as cytoskeleton of the body of the astrocytes (used in the AG index), stored in the neuropil (astrocytic elongations) or secreted to the ECM GFAP (used in the NG index) and produced in an uncontrolled manner by the glioma cells (glioma protein, used in the GP index) (Figure 2, B, C and D). GFAP pixels were manually sampled from ICH stained images and collected in a library of pixels. The values of each color channels, red, green and blue, were analysed for further segmentation by thresholding.

**Figure 2:**
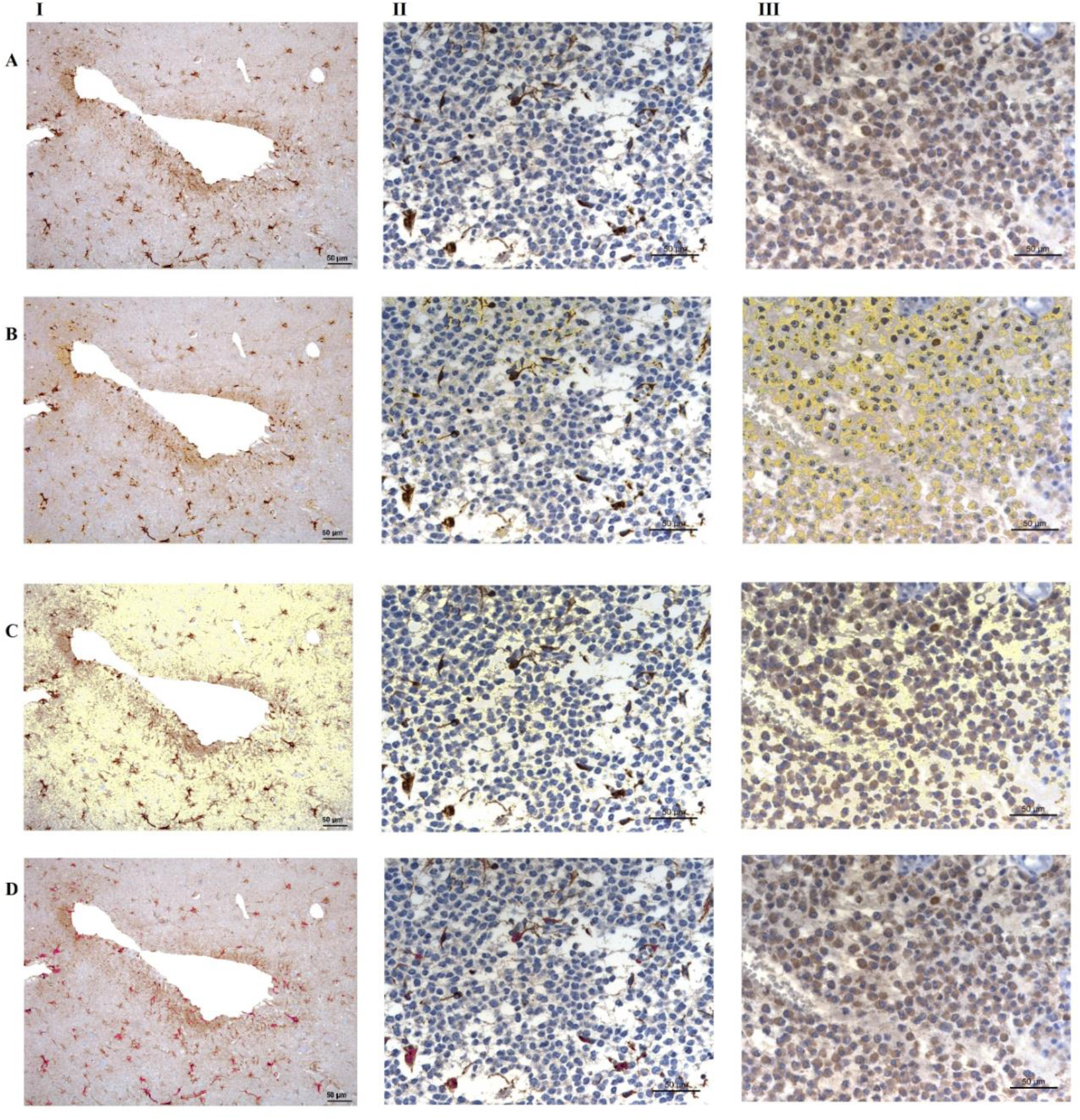
Brain tissue stained with GFAP immunohistochemical protocol. Columns: I: Normal healthy tissue. II: Stage III tumor. III: Stage IV tumor. Rows: A: Original GFAP image. Nuclei are stained in blue while the protein is brown. B: Protein segmentation of glioma GFAP in yellow overlapping original image. C: Protein segmentation of neuropil GFAP in yellow overlapping original image. D: Protein segmentation of astrocyte GFAP in pink overlapping original image.

Nuclei of the astroglial cells were stained with haematoxylin to differentiate from the target protein. The nuclei from the mutant astroglial cells were equally stained with haematoxylin but responded differently resulting in a darker color. A library of 865 pixels was also created for further segmentation.

For each structure *i* ∈ ℕ, 1 ≤ *i* ≤ 4,^1^ and each color channel *j* ∈ ℕ, 1 ≤ *j* ≤ 3, a coefficient *C*_*ij*_ was determined by essay-error, thus including the most certain values within the segmented feature (Table 2).

**Table 2:** Number of pixels measured for each feature, average value for color and coefficients for threshold segmentation.

| Structure | Pixels<br>sampled | Average |  |  | Coefficient |
| --- | --- | --- | --- | --- | --- |
|  |  | Red | Green | Blue |  |
| Astrocyte<br>GFAP | 967 | 90,13 | 19,84 | 9,65 | 1 |
| Neuropil<br>GFAP | 1515 | 210,94 | 199,53 | 202,06 | 1,5 |
| Glioma<br>GFAP | 1129 | 144,18 | 123,14 | 115,95 | 1,5 |
| Glioma<br>Nucleus | 864 | 104,55 | 103,03 | 128,48 | 2,5 |

These coefficients led to thresholding values defined by

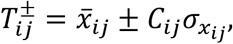

where 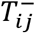 is the lower threshold value, 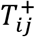 is the upper threshold value, 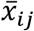 the average of the channel’s color value (between 0 and 255) over all pixels in the library for the given structure and 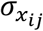 its standard deviation.

Given a pixel *x*, we define the thresholding function *ϕ*_*ij*_, acting on the *j* ^th^ color channel for the *i*^th^ structure by

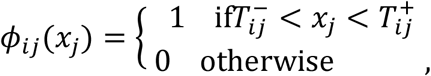

and the thresholding function *ϕ*_*i*_ for the *i*^th^ structure by

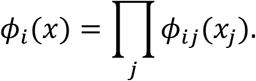

By applying each of the functions *ϕ*_*i*_ to all pixels in a given image, 4 binary images were obtained, one for each structure.

#### 2.5.3 Post-processing: Processing and quantification

##### 2.5.3.1 GFAP protein

Segmented images from the three proteins were smoothed with a line of 3 pixels length to denoise small artefacts not connected to the protein of interest. The number of pixels was included in a vector. Pixels in the image were recorded as area of reference.

##### 2.5.3.2 Glioma nucleus

The binary images obtained by thresholding the glioma nucleus were denoised using a disk of 1px size. Holes were filled with the function “imfill” and the connected nuclei were separated by watershed transform developed by Steve Eddins (Figure 3). Each of the connected areas in the final image was computed as one nucleus. Nuclei from the edges were deleted in alternative images. The number was included in a vector to estimate the nuclei number in each image and subset.

**Figure 3:**
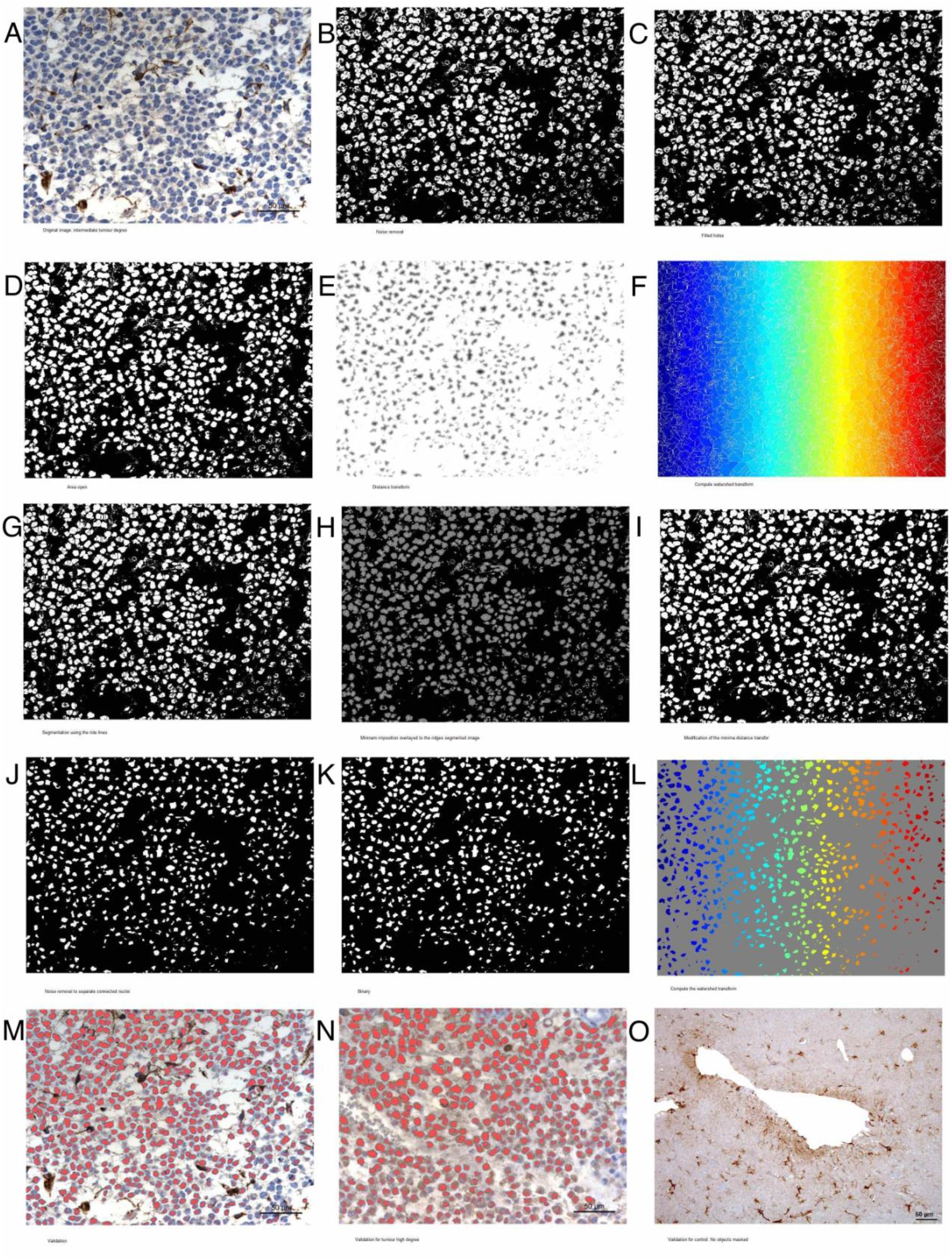
Watershed transform for glioma nucleus. A: Original image from stage III tumor grade. B: Noise removal. C: Filled holes. D: Area open. E: Distance transform. F: Compute watershed transform. G: Segmentation using the ride lines. H: Minima imposition overlapped to the rides of the segmented image. I: Modification of the minima distance transform. J: Noise removal to separate connected nuclei. K: Binary image. L: Computation of the watershed transform. Each one of the connected structures have been labelled and counted as one cell. M: Final validation of detected nuclei by overlapping in red on the original image of stage III tumor grade. N: Final validation in stage IV tumor grade. O: Application of glioma nuclei segmentation procedure to healthy tissue. No nuclei were identified thus evidencing the specificity of the segmentation for glioma nuclei.

### 2.6 Stereological indexes

The major features identified when tumors were graded by the pathologist were:

- GFAP found in neoplasic cells showing high density of protein and particularly placed around the nuclei (glioma protein henceforth). It was calculated as area and the units are μm^2^.
- Neuropil GFAP, corresponding to the lax GFAP found in the neoplastic intercellular space. As before, the units are μm^2^.
- Astrocyte GFAP, cytoskeleton found inside the body of astroglial cells with normal function measured by its area (μm^2^).
- Malignant nuclei (glioma nuclei henceforth). The units are nuclei number, or *nn*.

From these features, density ratios or indexes with physiological importance were developed:

- GP: the area of glioma protein per area of image (μm^2^/μm^2^) or no units
- NG: the area of neuropil per area of image (μm^2^/μm^2^) or no units
- AG: area of astrocytes per area of image (μm^2^/μm^2^) or no units
- GN: glioma nuclei number per area of image (nn/ μm^2^, number-density index)

The area of reference for density is the size of the image (μm^2^) and it varies depending on the analysis performed, being 86900μm^2^ for the complete image and 884μm^2^ for the subsets in the periodic and random sampling of the image.

Each image or subset that wasanalysed produced an array with GP, NG, AP and GN indexes to describe the presence of tumor features if any and its grade.

### 2.7 Precision of the image segmentation and processing

The precision of the segmentation by thresholding as proposed before was evaluated according to its sensitivity and specificity. To do so, an image of the segmented feature was overlapped with the original image to identify manually the segmentation (Figure 2B, C and D). The true positive segmented features (TP), the false positive (FP), the true negative (TN) and the false negative (FN) were counted manually in control, stage III and stage IV tumor histological images. The sensitivity was calculated as TP / (TP + FN) and the specificity as TN / (FP + TN).

Glioma and neuropil GFAP were evaluated by overlaying a grid of square cells on the image combined from the mask and the original. The true positives were those cells of the grid where the positive mask occupied more than the half of the protein that should be masked. The true negatives were the cells where non-protein areas were not masked, in case those areas were present. The false negatives were the cells where proteins that should be masked are not and the false positives were the cells where proteins that should not be counted are masked.

Astrocyte GFAP and glioma nuclei quantification were evaluated by overlaying the masked nuclei with the original image. Astrocyte GFAP true positives were those astrocytes with cytoskeletal GFAP providing star shape, typical from these cellules. True negatives were those cells with stained nuclei but corresponding to different cellular types that did not have any GFAP masked. False positives were GFAP proteins segmented that do not correspond with astrocytes and false negatives were astrocytes that have not been detected. In the case of the glioma nuclei, the true positives were the coincident nuclei. The true negatives were the healthy nuclei that were not masked as glioma. The false negatives were those glioma nuclei that were not detected. The false positives were those healthy nuclei or blood cells that were masked.

### 2.8 Multilevel statistics and characterization of the tumor

Tissue differences were characterized by a one-way ANOVA and multivariate analysis using R v. 3.1.0 GUI 1.64. The variables tested were the 4 variables defined previously, namely: GN, GP, NG and AG. The images used were the subsets created on the periodic image sampling (n=1415). As established by Weibel (1979), the variability between subsets has been taken into account as inter-individual variability random effect, and therefore, as source of error in the ANOVA model under an F-test. Further Pairwise T-test has been used to compare between tissues.

### 2.9 Classification for diagnosis: detection and grading

The two-step hierarchical classification was divided in detection (presence or absence of tumor) and further grading (grade III and grade IV) and the final classification was achieved by prediction after training.

Two training sets were constructed, one for tumor detection and another for tumor grading, including pathological cases in early or late stages and also inter-stage tumour grade accounting for inter-individual variability. First set for detection included the indexes from 30 subsets of control tissue (individual 10) and 30 subsets with tumor (individuals 1 and 2). The second set for grading included 30 subsets of tumor in stage III (individuals 1 and 9) and 30 subsets with tumor in stage IV (individuals 2 and 7).

PCA was performed in R for training the system and data was used for prediction in the classification model. Figures were obtained with Matlab and ggplot2 library from R.

Several general classification models were built: SVM, kNN and Gaussian decision tree (data not shown). According to the preliminary results the parametric Gaussian decision tree was the retained model since it fitted the best by its accuracy. Training was built up on a 10-fold loop CV and with the previously standardized data. Code for training function was generated with the Classification Learner app from MATLAB 2016a.

The classification used the Gini’s Diversity Index for split criterion in determining the nodes; the maximum number of splits was 4 for tumor detection (simple decision tree or SDT) and 20 for grading (medium decision tree or MDT). There were no surrogate decision splits.

The algorithm for classification was measured by use of the sensitivity (TP/FN) and positive predicted over false discovered (PP/FD). The model generated was used for further prediction with a trained algorithm.

The analysis pipeline started with the detection function followed by the grading function. The subsets were classified by the grading and each subset was then grouped according to the individual or patient (Table 1).

A test set for validation of the predictions was constructed with 1256 subsets from individuals 1 to 10. Those subsets that were used for training were excluded from the test set. The final report detailed the number of subsets without tumor, stage III and stage IV tumor found in each patient. A team of pathologists supervised the report to verify the final tumor detection and grading.

### 2.10 Software development

Image analysis was performed with MATLAB v. 2016a. For statistical analysis R c.3,1,0 GUI has been used. The code can be found in:

https://github.com/AylaScientist/GFAP-segmentation (segmentation)

https://github.com/AylaScientist/GFAP-classification (classification)

https://github.com/AylaScientist/GFAP-diagnosis (diagnosis)

## 3 Results

### 3.1 Histological and IHC evaluation

Tumors included in this study showed the typical pathological features of an Anaplastic Oligodendroglioma (WHO grade III) and Glioblastomas (WHO grade IV). According to their malignancy grade, glioblastomas showed a stage IV of cytonuclear atypia and an increased frequency of necrosis and glomeruloid vascular proliferation. Pseudo-palisading of neoplastic cells around necrotic foci was a pathognomic feature of Glioblastoma. Additional features observed were and infiltrative growth pattern, muccinous secretion, and some areas of inflammation.

### 3.2 Color differences between features from healthy tissue and glioma

One-way ANOVA applied to the pixel library has shown high significant differences for each of the three color channels RGB for all the tested structures, namely astrocyte, nucleus-high, GFAP control, GFAP-high (Supplemental 1, 2, 3 and 4, p value = 0, F-test). The intensity of the color is higher for the glioma nuclei while it is lighter for the healthy nuclei. Similarly, GFAP synthesized by the healthy astrocytes is significantly darker than the one synthesized by the mutated glial cells, which in turn are darker than the GFAP protein found in the neuropil (p value = 0, F-test and Tukey multi-comparison test).

### 3.3 Precision of GFAP image segmentation

The precision of the segmentation is described in Table 3. There was a notable decrease in the specificity of the glioma nuclei down to the 61% for tumors of stage IV. This decrease is due to a high number of false positive (46 false positive compared to 72 true negative) corresponding to artefacts and blood cells that were stained as nuclei but were not. However, no healthy nuclei were masked as glioma nuclei. Moreover, the segmentation did not detect any nuclei of the glioma in the normal tissue images.

**Table 3:** Accuracy of the segmentation of the tumor structures. Values are in percentage. The nucleus glioma couldn’t be evaluated in the healthy tissue because this structure was not found. The astrocyte GFAP couldn’t be evaluated in the stage IV tumor because no astrocyte remained in this tumor grade. The protein secreted by tumor cells or glioma GFAP can’t be evaluated in the control tissue since there is no tumor protein expressed.

|  |  | Healthy | Stage III | High |
| --- | --- | --- | --- | --- |
| Nucleus glioma | Sensitivity | - | 88.5 | 91.7 |
|  | Specificity | - | 88 | 61 |
| Glioma GFAP | Sensitivity | - | 100 | 100 |
|  | Specificity | - | 84.1 | 94.1 |
| Neuropil GFAP | Sensitivity | 75.6 | 88.6 | 79.1 |
|  | Specificity | 100 | 100 | 100 |
| Astrocyte GFAP | Sensitivity | 87.3 | 100 | - |
|  | Specificity | 94.88 | 94.59 | - |

The false negatives in the segmentation for protein secreted for neuropil (neuropil GFAP) correspond to very lax protein that was not masked in the healthy and stage IV tumor tissue.

### 3.4 Reduction of the standard error by sampling histological images

The use of subsets from complete histological images was applied here for reduction of standard error as suggested by Weibel (1979). The standard error was reduced for all the features below 5%, except for the astrocyte GFAP protein, which remains around 10% (Figure 4). The periodic sampling analysis show a significant reduction of the error when compared to the random sampling method.

**Figure 4:**
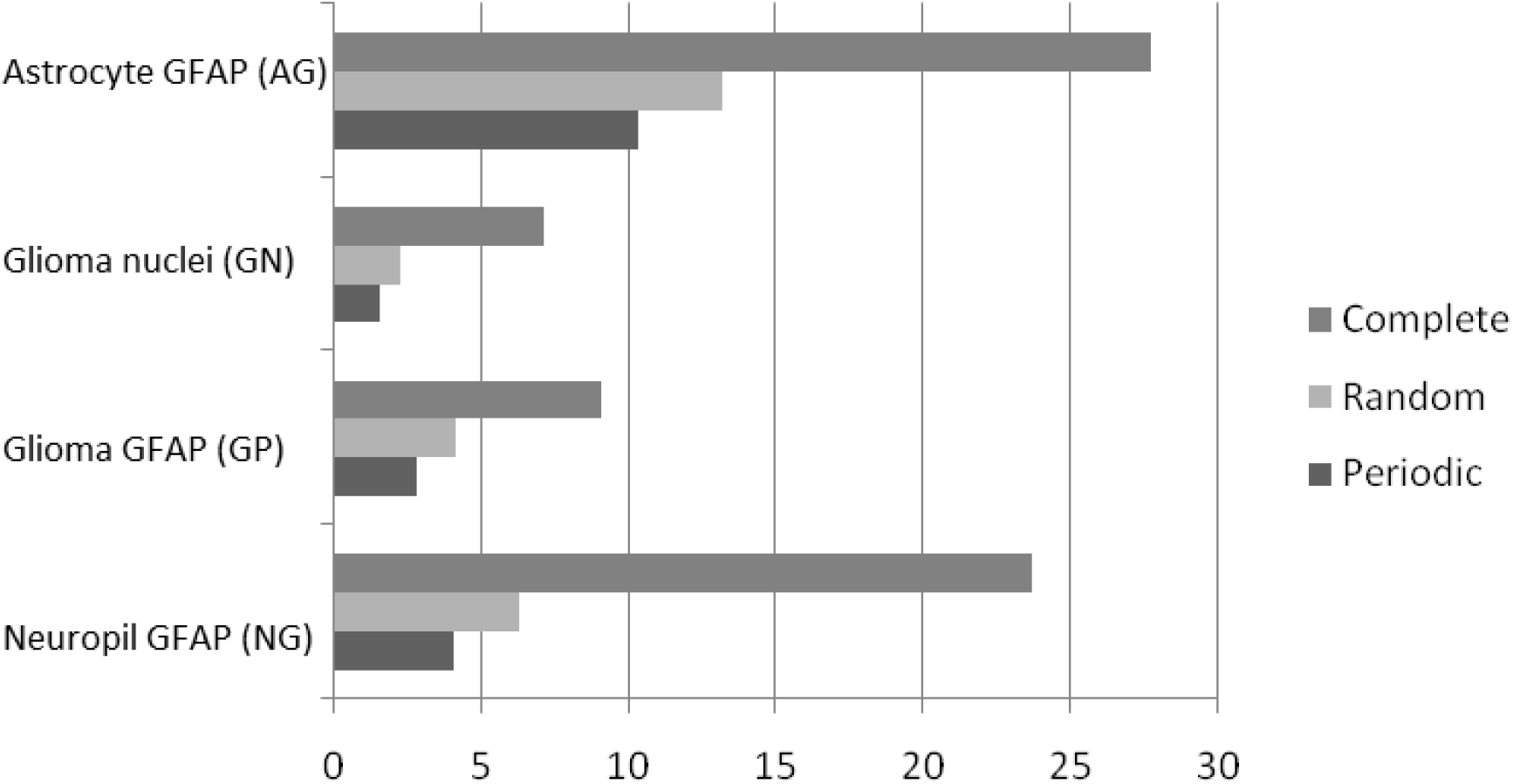
Standard error of the mean of the stereological indexes by each image sampling method: Complete, Random and Periodic. The error is expressed in percentage. The indexes are astrocyte GFAP density (AG), glioma nuclei number-density (GN), glioma GFAP density (GP) and neuropil GFAP density (NG).

### 3.5 Tissue differences detected by multilevel statistics

The presence of the tumor was easy detectable by the presence of glioma nuclei (GN), that did not exist in the normal tissue (Figure 5A, p-value = 0, F-test and T-test). Similarly, the glioma GFAP (GP) was significantly higher in the glioma tissue (Figure 5B, p-value = 0, F-test and T-test). Both GP and GN were significantly higher in tissues with stage III tumor grade when compared to stage IV (Figure 5B, p-value = 0, T-test).

**Figure 5:**
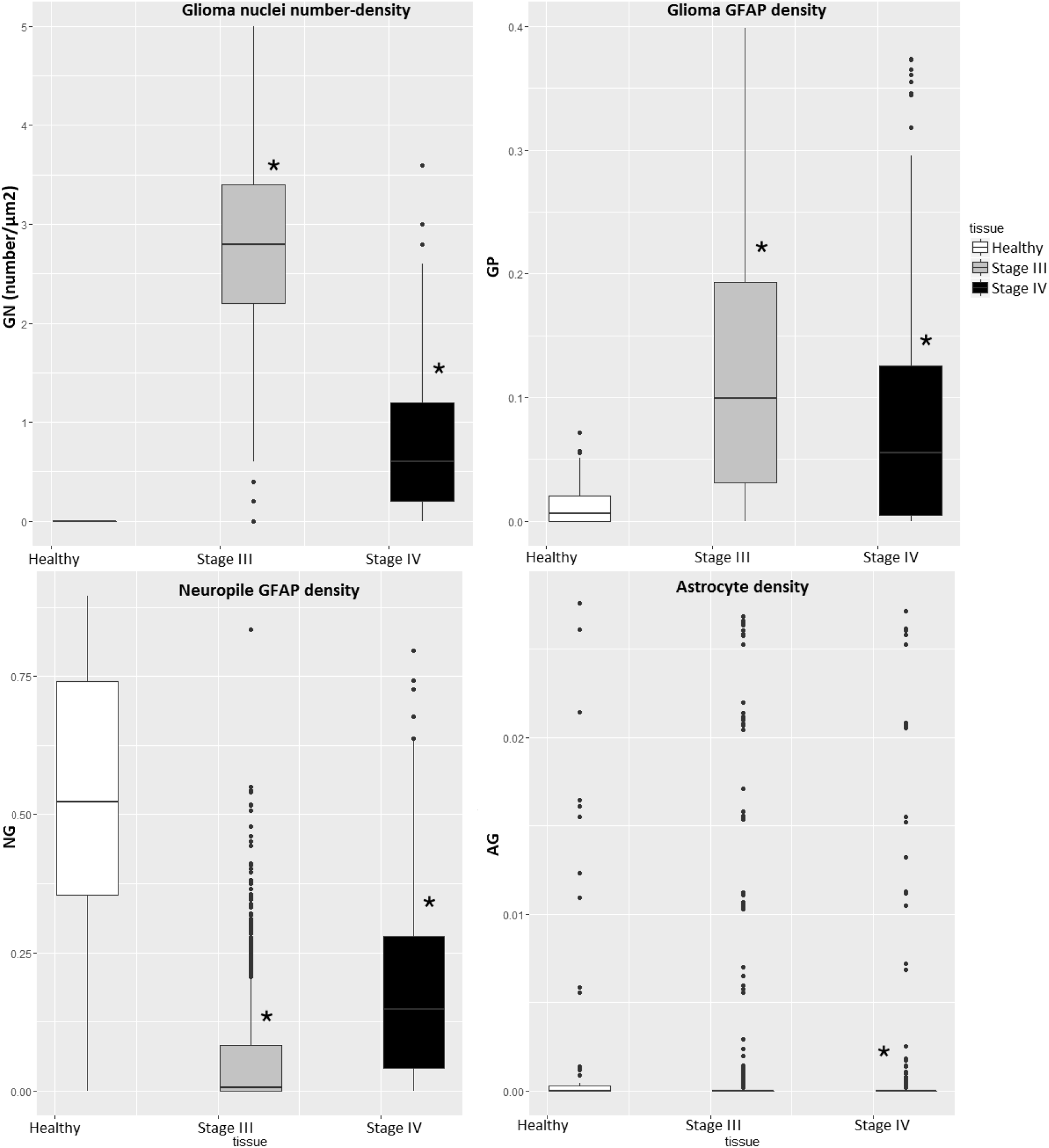
Multilevel statistics applied to GFAP quantification. A: Glioma nuclei number-density (GN). B: Glioma GFAP protein density (GP). C: Neuropil GFAP protein density (NG). D: Astrocyte GFAP protein density (AG). Significant differences marked with *, p-value = 0 for both F-test and T-test.

NG was significantly higher in normal tissue than in the tumor samples. It was also higher in stage IV tumor when compared with the one in stage III (Figure 5C, F-test, p-value = 0).

AG was significantly higher in the healthy brain tissue and in the stage III tumor when compared to the stage IV tumor (Figure 5D, F-test, p-value =0). However, the difference between stage III tumor and normal brain tissue was not significant due to the low density of astrocytes in both tissues and the high dispersion of this index.

### 3.6 Classification and tumor diagnosis

The best algorithm for classification is the SDT for tumor detection and MDT is the best for tumor grading according to their accuracy (Table 4).

**Table 4:** Accuracy for tumor detection and grading used in the diagnosis pipeline. TP/FN: total positives over false negatives. PP/FD: predicted positives over false discovered.

| Phase of diagnosis | Training algorithm | Accuracy Matlab application | TP/FN (%) |  | PP/FD (%) |  |
| --- | --- | --- | --- | --- | --- | --- |
|  |  |  | Healthy | Tumor | Healthy | Tumour |
| Detection | Decision tree | 93.3 | 97 | 90 | 91 | 96 |
|  |  |  | Stage III | Stage IV | Stage III | Stage IV |
| Grading | Decision tree | 86.7 | 90 | 83 | 84 | 89 |

The pipeline for diagnosis trained with the first set of samples provided low percentage of sets in the correct diagnosis for high stages tumors and in particular for individual 7, thus classifying it as stage III or even healthy. However, the second training set provided a majority of subsets classified in accordance with the pathologist criteria for all pipelines (Table 5).

**Table 5:** Results of the PCA showing the importance of the neuropil (NG), the glioma nuclei density (GN) and the glioma GFAP (GP) in the first component. The second component is characterized mostly by the astrocyte GFAP (AG) and secondarily by the glioma nuclei density (GN). The third component is affected mostly by the glioma GFAP (GP) and the fourth component again by the neuropil and the glioma nuclei density. The first component explains the 44.85% of the variance, the second is 25.44%, the third is 18.52% and the fourth is 11.9%.

| Standard deviations |  |  |  |  |
| --- | --- | --- | --- | --- |
|  | 1.3393724 | 1.0087375 | 0.8607095 | 0.6691111 |
| Rotation |  |  |  |  |
|  | PC1 | PC2 | PC3 | PC4 |
| NG | -0.63335577 | 0.06689965 | 0.1821262 | 0.7491428 |
| GN | 0.57626118 | 0.21565878 | -0.5183099 | 0.5939437 |
| GP | 0.51093318 | -0.01598525 | 0.8275075 | 0.2322135 |
| AG | 0.07570209 | -0.97404322 | -0.1158283 | 0.1791446 |
| Importance of components |  |  |  |  |
|  | PC1 | PC2 | PC3 | PC4 |
| Standard deviation | 1.3394 | 1.0087 | 0.8607 | 0.6691 |
| Proportion of Variance | 0.4485 | 0.2544 | 0.1852 | 0.1119 |
| Cumulative Proportion | 0.4485 | 0.7029 | 0.8881 | 1.0000 |

### 3.7 PCA

The PCA shows a strong correlation between NG, GN and GP indexes that contribute to the first principal component explaining 44.85% of the population’s total variance (Figure 6, Table 5). GN and GP indexes contribute positively to this component, whereas NG contributes negatively. It shows that the relation of NG is positively correlated to the health status of the brain tissue, while GP and GN overlap the different tumor stages. The second principal component is mostly composed by AG, with a negative contribution and explains 25.44% of the total variance. GN and GP are the two components mainly contributing to the third principal component, explaining 18.52% of the total variance. Finally, the fourth principal component that explains 11.19% of the population’s variance is a mix of all original components, dominated by NG and GN.

**Figure 6:**
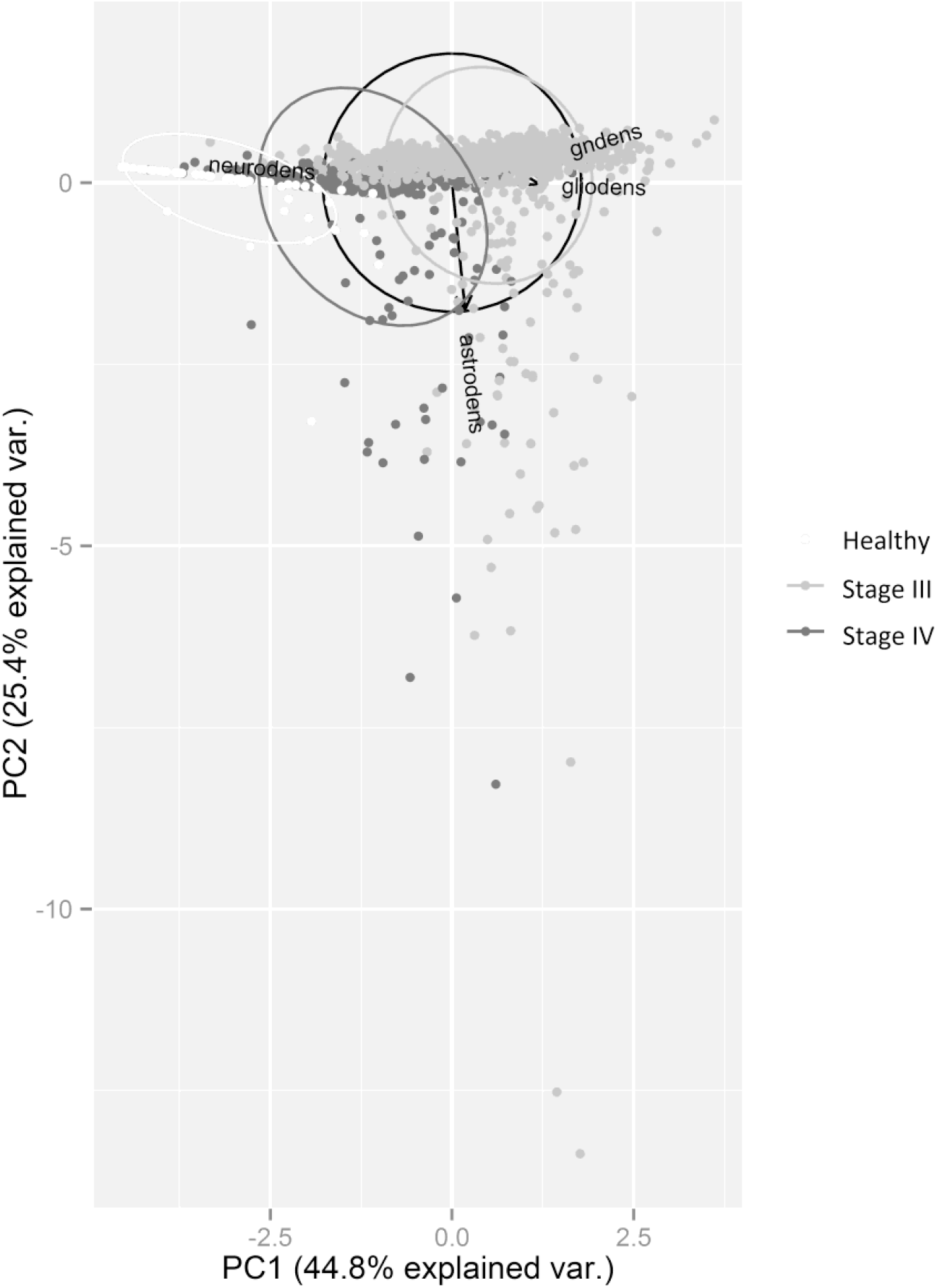
The data plotted in the frame of the two first components of the PCA that comprise 70.29% of the total variance. Neurodens corresponds to neuropil GFAP density (NG), gndens to glioma nucleus number-density (GN), gliodens to glioma GFAP (GP) and astrodens to astrocyte GFAP density (AG). NG, GN and GP have major contribution to the first component, thus establishing a linear progression. Scaling is 1. Ellipses include the 95% of the samples from each tissue marked by the color and the centre is obtained by the mean of the group. Circle in brown has 2 units radius and is centred in the point 0,0. Graph is obtained by with the package ggbiplot in R.

The 4 principal components were used in the learning process to cover the maximum of the population variance. This choice was later supported by the results of the different training sessions of the classifier.

## 4 Discussion

### 4.1 Canine glioma as human model

Previous studies have demonstrated that canine and human gliomas of different cell lineage and malignancy grades share similar histopathological and IHC features [39] as well as molecular expression profile [40,41] and biological behaviour [42]. For rodent gliomas, the histopathological features do not always agree with observations in their human counterparts and prediction of therapeutic response is poor [43,44]. These data, together with the equivalences in terms of tumor initiation and progression [45,46] support the use of canine gliomas as model of human gliomas.

The results obtained in the present study provide insights for the understanding of glioma development and cancer biology.

### 4.2 Quality of the image analysis

In the brain, intermediate filaments of astrocytes include the GFAP form the cytoskeleton of the astroglial in the central nervous system CNS. This protein is also found at a lower density in the extracellular matrix (ECM) and in the elongations of the astrocytes in the neuropil. The neuropil is a synaptically rich region of the brain composed by dendrites, unmyelinated axons and glial arborisations. It forms the bulk of the central nervous system’s grey matter and in which the neuronal cell bodies lie embedded. The function of the neuropil varies, depending on the area of the brain, and it is mostly related with the required synaptic connections of the zone. The alteration of this protein during oncological processes has been described widely [47–49] and also for dogs [39,42,50]. Therefore, accurate selection and segmentation of the GFAP for glioma diagnosis is one of the key factors for the success of the grading method validated here.

The stereological approach by subsetting the image and developing four unbiased indexes of reference increased the accuracy of the present pipeline. Subsequent classification by a trained system is the final step on this diagnosis procedure. Inter-individual variability has been taken into account to train the system as a necessary approach. In addition, this machine learning study integrates the inter-individual variability along the tumor phases as a source of error in the classification.

However, the procedure can be improved to detect palisades or pseudopalisades by differential stain of the cancer nuclei using proliferation markers such as Ki-67 or cyclin A.

### 4.3 Identification of key structures for grading

A board of pathologists assessed the histological evaluation of the tumors with IHC specific for GFAP, following the WHO directives [38]. The features used during this analysis have been successfully segmented with high levels of precision (Figure 5) thanks partially to the specific staining of the GFAP.

Additionally, the successful identification of three different proteomic patterns that attach to the same antibodies constitutes a novel approach for histological diagnosis. The quantity of antigen and colour attached to the protein during the staining procedure depends on the density and the conformation of the GFAP in each feature of the brain or glioma tissue. The Lambert-Beer law [51] or light absorption determines the absorbency of a given wave length in a sample depending on the quantity of absorbent species that the light finds when passing through the sample. Since the agglomeration of the normal astrocyte GFAP includes higher density of protein than the glioma, there is also more antibodies attached, thus coloring it significantly darker. That also applies to the neuropil GFAP in the contrary case: the neuropil has a very lax GFAP and therefore the color is significantly lighter. This biophysical property allows the differential segmentation of the three different protein conformations.

The protein produced in the glioma cells is agglomerated in a different manner than the healthy cytoskeletal one synthesized by the healthy cells. This difference can be explained by the high mitotic activity of the tumor cells [31]. The tumor cell physiology is affected in a manner that cannot process the GFAP thus losing the shape and as proposed by Chen and co-workers [52], also changing the function.

The same principle applies to the differential staining of the cancer cell nuclei when compared to the health cell nuclei. Again, because of the high mitotic activity described by Gichet [31], the staining of the cancer nuclei has higher accumulation of colorant, and consequently, a darker color. Therefore, the differential segmentation is possible because of the differential pattern of the chromatin structure.

### 4.4 Segmentation

The sensitivity and specificity of the GFAP structures is maintained over 84%, with an exception of the sensitivity of neuropil GFAP that decrease to 75% in healthy tissue and 79.1% in stage IV tumor. The library of pixels for segmenting the neuropil was created from the more identifiable color in the neuropil of the healthy tissue, which is also the densest neuropil GFAP. This GFAP density is also similar to the one found in the neuropil of a tumor. However, the very lax GFAP in the neuropil of healthy tissue is not segmented since it has very similar coloration than the background. Those different densities of the protein associated to neuropil can be influenced by the ECM, which can also contain almost degraded GFAP from cellular processes. That can be explained by the heterogeneity of the ECM in the normal brain tissue and also during tumor development [53]. Nevertheless, the area of the ECM in brain tissue is considered almost non-existing and the specificity was 100%. Therefore, the reduction in the sensitivity for the neuropil did not reduce the accuracy of the segmentation. That is also consistent with the high accuracy of the further tumor classification. This method opens here for the first time the possibility to quantify and to study the heterogeneity of the brain neuropil with unbiased quantitative tools.

The accuracy values are all above 88% for nuclei segmentation with the exception of the nuclei in the stage IV glioma (IV), showing 61% of specificity. This can be explained by the increase of the necrosis areas and capillaries in the advanced stages of the tumor [38]. The dead cells and the blood cells in the tissue are also colored as nuclei and therefore the segmentation process has false positives. This problem could be possibly solved with differential stain for proliferation such as Ki-67 or cyclin A [4]. However, the sensitivity of the segmentation here was enough to compensate the low specificity for the tumor in grade IV (91.7%) and the accuracy was not reduced by the presence of false positives. Even though this reduction in accuracy did not affect the final diagnosis of the tumor, a complementary evaluation of the vascular components, such as capillary glomeruloid, can be recommended.

The identification of the nuclei by its color and its separation with watershed techniques has been also applied before by Mouelhi [6]. However, these studies focused only on the density of nuclei as markers and are not strong enough to develop a proper diagnosis. In our study, the watershed segmentation applied to the tumor nuclei is a complement to the GFAP indexes thus creating a complete array of cancer indexes.

### 4.5 Stereological indexes

Subsetting the image by either periodic or random sampling reduced the standard error of the indexes when compared to the analysis of the entire image. Furthermore, the periodic sampling is proved to be more accurate because it has a higher number of subsets, and therefore, the indexation is more robust. The standard error of the indexes is reduced below 5% in all the cases except in the astrocyte-GFAP density that remained in the 10.33%. This exception occurs because the distribution pattern of the astrocytes has a frequency of appearance that is bigger than the period of the sampling. It means that only few subsets have astrocyte-GFAP. The most subsets in all the sampled tissues have no presence of astrocyte-GFAP and it produces a binomial distribution of the frequency: or no astrocyte (majority of subsets) or certain density, depending on the subset (minority). However, the astrocyte GFAP has been proved to be a determinant factor for grading of tumour [38]. Therefore, this index has been also included in the classification-learning algorithm with a particular success in grading.

Furthermore, the neuropil is providing here a new pattern for glioma diagnosis. The total area of neuropil is significantly reduced by the cell number increase during the glioma development. Added to it, the records on accuracy for neuropil segmentation have detected false negatives within the healthy brain tissue due to a very lax protein that could not be segmented. These results are indicating an increase in the protein density and possibly the rigidity of the synaptic area and ECM during tumor development, in agreement with typical cancer growth patterns [54]. This variation may change the contact area between cells and cell-to-ECM, altering the cell activity, communication and signal spreading through the tissue. By quantifying the neuropil during tumor development as proposed here it is possible to approach the synaptic changes, the cell activity and the mechanical deformation of the cell. Moreover, changes in the cell binding and in the ECM could modulate the cell growth and viability [52], and determine tumor development. Actually, the recently described glioneuronal tumors with neuropil-like islands have been not surprisingly identified as molecular astrocytomas [55], thus illustrating the alterations that the neuropil can suffer during tumor growth. Therefore, the quantification of the neuropil is revealed here as a major factor in tumor detection and grading, as important as tumor cell density. Indexes depending on the protein content and conformation of the neuropil will be a tool for research in aggregates of Huntington’s Disease [56] and neuropil threads of Alzheimer [57], among many other fields in neurophysiology.

The cytoskeletal structure of glioma GFAP has shown also to be a useful indicator of the tumor presence, cell lineage origin and grade of the tumor, in agreement with previous studies [58]. The abundance of this protein conformation is gradually increasing along the tumor development till arriving to stage III. However, the density of glioma cytoskeletal protein is reduced on stage IV when compared to stage III. This can be related with a higher grade of undifferentiating neoplastic cells after stage III. That agrees with previous reports of less differentiated glial precursor cells involved in the poor clinical outcome of high grade gliomas [59,60].

The cancer nuclei number-density has been used mostly in studies as a main indicator for tumor detection and grading [4,6,17,38]. Other scoring methods for cancer such as nuclear and membrane markers include counting the percentage of positively stained nuclei or area (pixels) to determine the presence of a tumor [19]. However, the creation of indexes with a constant area of reference provides here the accuracy of the unbiased quantification for further comparative analysis [29,61]. Added to it, nuclear markers for proliferation such as Ki-67 or cyclin A [34,62] could be used as a complementary approach to the conformation of glioma GFAP for prediction of palisades and pseudopalisades typical of the stage IV.

The stereological indexes developed here shown a very low standard error and a high accuracy for tumor diagnosis. All of the four indexes developed have an important role in tumor detection. Added to it, the density of astroglial protein is significantly reduced only in the tumor grade IV, and therefore, the AG index has a major role in supporting the grading and a minor use for detection.

### 4.6 PCA: interpretation of the results

The PCA illustrate the importance of all the indexes in the diagnosis. The most important index in the first component is the protein in the neuropil closely followed by glioma protein and nuclei. These indexes point the evolution of the tumor mediated by the neuropil. These results reveal a pattern: the higher the density of tumor cells, the lower the density of neuropil. This pattern may also indicate a reduction or modification of the synaptic activity within the tumor brain tissue.

Two indexes, GP and GN, are contributing negatively to the first component. This is an indicative of the production of glioma GFAP by the high activity of the glioma nuclei. Therefore, this first component includes the necessary variance for tumor detection. Furthermore, the astroglial GFAP is the most important index contributing to the second component. The reduction of AG density is indicative of tumor presence, in agreement with the WHO classification of tumors [38]. Therefore, this second component shows a particular importance for tumor grading, illustrated by the astroglial protein. AG is also counteracted by the presence of glioma nuclei, thus indicating the importance of the healthy cell substitution by malignant ones in the development of the tumor.

The third component is represented by GP and GN, thus indicating the presence of a tumor. The fourth component explains the relation between the reduction of the neuropil and the increase of the glioma nuclei. This linear relation indicates the tissue turnover during the tumor growth, which in turn is also an indicator of tumor grade.

### 4.7 Classification learner

The machine-learning process for classification of tumor is proved more efficient for tumor detection than for grading (Table 4). That responds to the clear binomial patterns for tumor, such as presence or absence of glioma nuclei or glioma GFAP. In our tumors, the progressive reduction of AG and glioma nuclei when collapsing the tumor in the grade IV allowed an efficient tumor malignancy grade prediction, according with the inverse relation with the expression of GFAP attributed to human gliomas [63–65]. Added to it, the changes in the neuropil are also indicative of each stage and therefore of importance for both detection and grading.

The classification learner has been evaluated in terms of percentage of correctly classified subsets. At least half of the subsets from the same individual must accomplish the same criteria than the pathologist in order to be valid. This difference of criteria for subsets belonging to the same individual is in accordance with the microareas of variability present in all the tissues [66]. These areas can be captured here thanks to the image subsetting. When working with a tissue in constant transformation, as is the development of a tumor there are microareas with full tumor development and healthy ones [38]. Consequently, it is desirable to reflect this variability in the classification of the subsets.

The most challenging diagnosis is the one for tumors inter-grade such as stage III and stage IV. These differences cannot be recognized if the training set establishes strict patterns for grade standards. Because of the continuous development of the tumor, classification methods should also recognize inter-grade tumors to select a proper treatment. This was mostly represented by the individual number 7, which corresponds to an early stage IV tumor. The tumor of this individual shows higher amount of cancer nuclei when compared with a theoretical model of stage IV glioma as for example individual 2. Individual 7 was considered healthy or grade III patient indistinctly when the system was trained without representing the early stage IV. Only after training the system with a more heterogeneous set, the diagnosis can be corrected for all patients. This is also an indication that a training set collecting only the patterns for each tumor grade as established in the WHO classification cannot always detect the inter-grade tumors.

The most critical medical decision making for treatment selection corresponds to grades III and IV, the ones successfully classified here. However, a medical research approach should include also grades I and II in order to achieve a full perspective of the tumor development.

## 5 Conclusions and recommendations

The role of the neuropil in the development of gliomas is quantified here and compared to those of the tumor cell proliferation and growth. The density of neuropil GFAP protein is the most important index for brain tumor diagnosis according to the PCA. This is a new approach for brain histological evaluations. Unbiased quantification of the neuropil can be included in order to approach cell communication in the physiology of the whole tissue, thus opening new research horizons within the neuropil field.

This is the first time that a whole pipeline of accurate segmentation integrates a successful machine learning approach for brain tumor diagnosis. When compared with other stereology software, the application developed here successfully automates the quantification and classification of both tumor and healthy brain tissue, thus avoiding time for manual counting. The four indexes developed here constitute an array able to determine the presence of tumor and also the possibility to grade its malignancy in a successful diagnosis process.

As a novel approach, we discovered here that the IHC stained protein can have many different proteomic states depending on the tissue structure and developmental stage which in turn increases the efficiency in automated medical diagnosis. A suggestion for further histological quantifications is to develop IHC and create a library of pixels in accordance to automate the diagnosis processes.

This IHC technique has also proved to be necessary for the classification under a machine learning approach. This is the first time proved that inter-individual variability for the tumor development must be classified with a less accurate training set. This variability should be taken into account henceforth when developing classification and grading of highly “transformable” tissues, such as tumors. This is a basic approach for achieving a correct diagnosis and prognosis, as well as for treatment selection with automatic semi-supervised machine-learning tools.

Finally, after increasing the efficiency in the indexation with stereology, the realistic inter-individual variability included in the training set allowed a successful detection and grading of the tumor for all the pathological cases exposed here.

## Acknowledgements

We gratefully acknowledge professor Antonella Zanna from the University of Bergen and MOLIMAGLIO (SAF2014-52332-R) for supporting this study. This research did not receive any specific grant from funding agencies in the public, commercial, or not-for-profit sectors. Author AC held a EU-ITN grant from the European Commission within Horizon2020.

## Footnotes

1 Where 1 is astroglial GFAP, 2 is neuropil GFAP, 3 is glioma GFAP and 4 is glioma nuclei.

